# Genome-Wide Mapping of 5’ Isoforms with 5’-Seq

**DOI:** 10.1101/2022.10.26.513882

**Authors:** Zlata Gvozdenov

**Affiliations:** Harvard Medical School, Department of Biological Chemistry and Molecular Pharmacology, 240 Longwood Ave, Boston, MA 02115

**Author notes:** address correspondence to: Dr. Zlata Gvozdenov.

**Keywords:** transcription start site, 5’ transcript isoforms, transcriptome, RNA sequencing, 5’-cap

## Abstract

Transcriptome of a genome is appreciated to be more complex than previously assumed. Same gene readouts can differ in terms of transcription start site, transcription end site and splicing. Growing evidence suggests functional importance of distinct transcript isoforms of the same gene. Obtaining these isoforms easily experimentally and processing data is crucial for prompt transcriptome functional characterizations. Here, I describe a quick protocol for generation of capped 5’ isoforms sequencing library and 5’ isoforms data analysis. The protocol relies on utilization of dephosphorylation-decapping method (oligo-capping), and it is a simplification of previously published 5’ isoform studies. The pipeline for data analysis suggests several isoform features to focus on.

## INTRODUCTION

Transcription initiation begins when an RNA polymerase binds promoter DNA and synthesizes the first nucleotide for an RNA molecule. The start of an mRNA molecule in eukaryotes is characterized by the presence of a chemical modification, 5’-cap (Shatkin, 1976). The DNA sites where transcription initiates, usually in 5’ untranslated region (UTR) upstream of a start codon, can vary leading to production of different 5’ isoforms of the same genes (Carninci et al., 2006). Utilization of alternative transcription start site as a mode of gene regulation was shown to be relevant for several biological processes (Davuluri et al., 2008, Zhang et al., 2017, Reyes and Huber, 2018, Ray et al., 2020) demonstrating a need to understand 5’ site selection, 5’ site divergence and its functional consequences. Crucial regulatory sequence motifs encoded within 5’ UTRs (present in longer and absent in shorter 5’ isoforms) could be important for the differential biological outcomes. Deriving such novel sequence elements prerequisites determining precise location and abundance of single 5’ isoforms on a genome-wide scale. This protocol describes experimental and bioinformatical pipelines to identify and quantify relative abundance of genome-wide 5’ isoforms of capped mRNAs, and it is also suitable for studying low intensity 5’ isoforms. The procedure can be applied to eukaryotic systems to shed light onto the functional importance of 5’ isoform heterogeneity *in vivo*.

Several experimental approaches for studying 5’ isoforms are available (Maruyama and Sugano, 1994, Carninci et al., 1996, Zhu et al., 2001, Scotto-Lavino et al., 2006, Pelechano et al., 2014, Machida and Lin, 2014, Pelechano et al., 2016, Jiang et al., 2019), each of which has its pros and cons, as reviewed (Policastro and Zentner, 2021). Here, I describe a simpler, faster, and relatively cheap procedure for the construction of capped 5’ isoform sequencing library, as outlined in Fig. 1. The procedure begins with enrichment and fragmentation of mRNA. The protocol is further based on dephosphorylation of mRNA molecules followed by decapping of 5’ mRNAs to leave 5’ monophosphate ends. This allows for the ligation of the sequencing adapter to the previously phosphorylated 5’ mRNAs (aka oligo-capping). 3’ adapters are ligated to 3’ RNA ends, the template is reverse transcribed and amplified for the high-throughput sequencing. The protocol utilizes advantages of magnesium-based RNA fragmentation, which does not introduce new phosphorylated 5’ ends, and can be performed before 5’ ligation to generate sequencing library compatible, smaller RNA fragments. The protocol significantly simplifies the procedure by relying on similarity of reaction conditions and heat inactivatable enzymes to perform a considerable number of crucial enzymatic reactions in the same tube successively. Besides time and cost reduction, the procedure avoids generation of intermediates via random oligos – the adapters are ligated directly on RNA from which the sequencing library is amplified.

**Figure 1.**
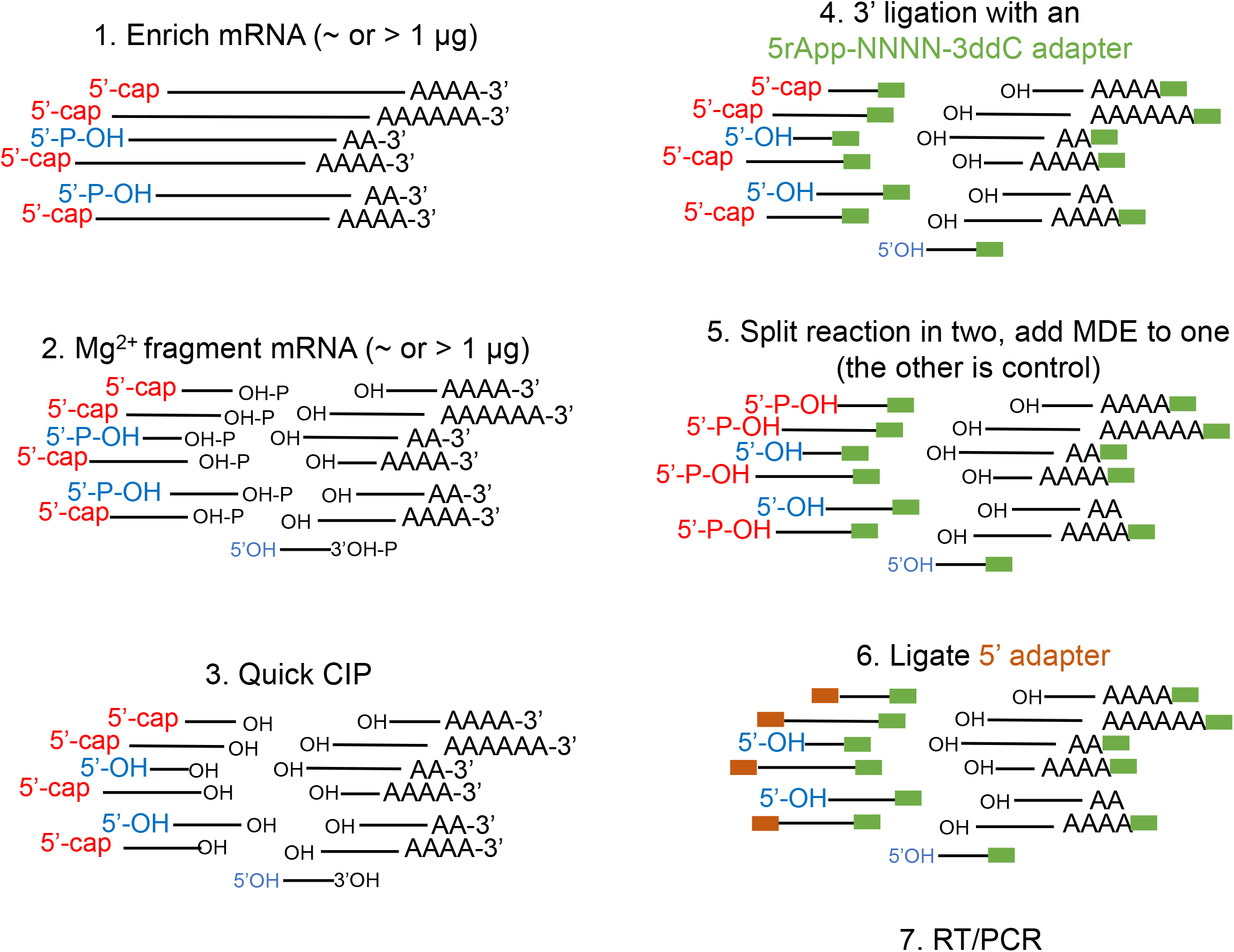
Overview of the 5’-Seq procedure. Messenger RNA molecules are enriched with poly(d)T beads and fragmented with magnesium module. The RNA fragments are dephosphorylated at both 5’ and 3’ termini and 3’ adapters are ligated. After this step, reaction is split into two tubes and decapping with mRNA decapping enzyme (MDE) is performed for one half (MDE+) while the other half of the sample is control (MDE-). 5’ adapters are ligated to decapped and phosphorylated RNA fragments, the fragments are reverse transcribed, and the library is PCR amplified for both treated (MDE+) and control (MDE-) samples.

## PROCEDURE FOR CONSTRUCTING SEQUENCING LIBRARY FROM CAPPED 5’ ISOFORMS

This section describes procedures for generation of indexed DNA library for high-throughput sequencing 5’ isoforms or transcription start sites using Illumina platform. This protocol is modified from other methods that utilize dephosphorylation-decapping tactics to study 5’ transcript ends (Tsuchihara et al., 2009, Ni et al, 2010, Gu et al., 2012, Arribere and Gilbert, 2013, Machida and Lin, 2014, Pelechano et al., 2014, Pelechano et al., 2016). DNA library purification of this protocol is comparable to earlier methods (Jin et al., 2015).

### Materials

100 μg purified total RNA, DNaseI treated

Nuclease-free water

Oligo d(T)25 magnetic beads (NEB, cat. no. S1419S)

NEBNext^®^ Magnesium RNA Fragmentation Module (10× fragmentation buffer and 10× fragmentation stop buffer, NEB, cat. no. E6150S)

3 M sodium acetate, pH 5.5

Ethanol (molecular biology grade)

15 mg/ml GlycoBlue (Thermo Fisher Scientific, cat. no. AM9516)

Quick CIP (NEB, 5 units/μl, cat. no. M0525S)

20 U/μl SUPERase.In (Thermo Fisher Scientific, cat. no. AM2694)

Oligonnucleotides (Table 1, also in Jin et al., 2015)

**Table 1.**
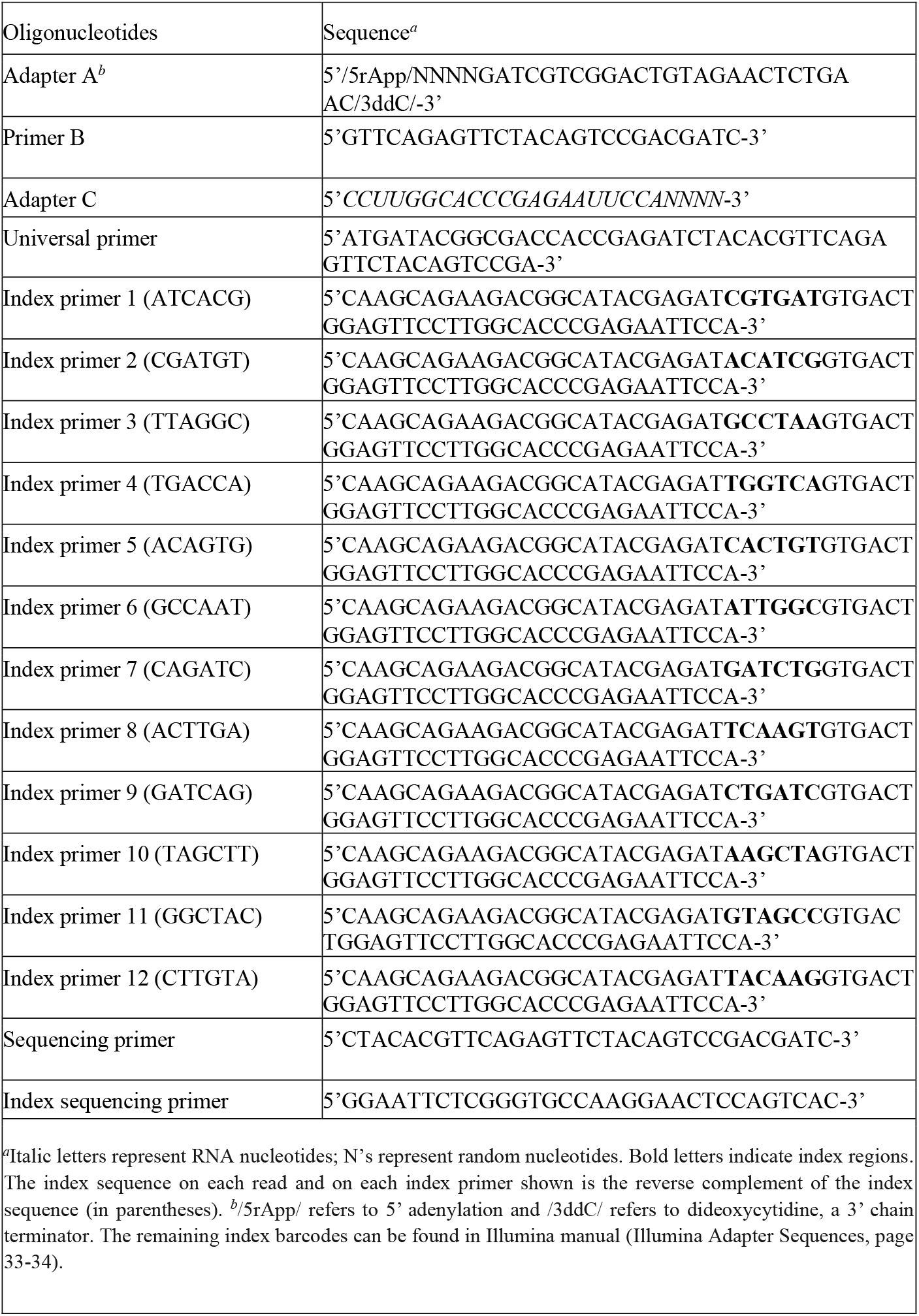
Oligonucleotide sequences (similarly to Jin et al., 2015).

30 U/μl T4 RNA ligase 1 concentrated (NEB, cat. no. M0437M)

10× T4 RNA ligase buffer (supplied with T4 RNA ligase 1)

10 mM ATP (supplied with T4 RNA ligase 1)

50% PEG8000 (supplied with T4 RNA ligase 1)

mRNA decapping enzyme, MDE (NEB, 100 units/μl, cat. no. M0608S)

200 U/μl truncated T4 RNA ligase 2 (NEB, cat. no. M0242L)

200 U/μl SuperScript III reverse transcriptase (Invitrogen, cat. no. 18080044)

5× First-Strand buffer (supplied with SuperScript III reverse transcriptase)

100 mM DTT (supplied with SuperScript III reverse transcriptase)

10 mM dNTPs

2 U/μl Phusion HF DNA polymerase (NEB, cat. no. M0530L)

5× HF buffer (supplied with Phusion HF DNA polymerase)

DNA SizeSelector-I (Aline, cat. no. Z-6001-50)

High-sensitivity DNA Kit (Agilent, 5067-4626)

6× Gel Loading Dye, Purple (NEB, cat. no. B7025S)

100 bp DNA Ladder (NEB, cat. no. N3231L)

PCR Marker (NEB, cat. no. N3234L)

TBE buffer (VWR, E442-500ML)

8% TBE polyacrylamide gel (Invitrogen, cat. no. EC62152BOX)

10,000× SYBR Gold Nucleic Acid Gel Stain (Invitrogen, cat. no. S11494) Ice

Liquid nitrogen or dry ice

Ice bucket

Safe-Lock microcentrifuge tubes (Eppendorf, cat. no. 22363204)

0.5-ml thin-walled PCR tubes

Magnetic stand

Shaking platform

Thermal cycler

25°C, 37°C, 50°C, 55°C, 65°C, 70°C, 80°C, 94°C heating blocks

Refrigerated tabletop centrifuge

Vortex

Gel electrophoresis apparatus

2100 Bioanalyzer system (Agilent)

Disposable scalpel

Transilluminator

20-gauge needles

Costar Spin-X centrifuge tube filter (0.45-μm cellulose acetate in 2-ml tube; Corning, cat. no. 8161)

### Preparation of beads for mRNA purification

Prepare oligo d(T)25 magnetic beads for use in step 6.

1. Equilibrate the oligo d(T)25 magnetic beads to room temperature (RT) for 20 min prior to use. *Before taking an aliquot, mix oligo d(T)25 magnetic beads thoroughly by inverting and/or swiveling the bottle.*
2. Wash 200 μl of bead slurry with 1 ml of 1× annealing buffer in a 1.5-ml microcentrifuge tube (per RNA sample; scale appropriately). Collect the beads on a magnetic stand and discard the supernatant. Repeat the wash one more time. *Do not wash more than 500 μl of original bead slurry with 1 ml buffer (use additional 1.5-ml microcentrifuge tubes).*
3. Resuspend the washed beads in 400 μl of 1× annealing buffer per RNA sample.

### Isolation of mRNA

4. Dilute 100 μg of high-quality total RNA with 400 μl of nuclease-free H_2_O in a 1.5-ml microcentrifuge tube before adding 400 μl of 2× annealing buffer. Split reaction in 4 tubes (200 μl each). *RNA can be prepared by either a hot-phenol extraction method or a TRIzol extraction method followed by purification using RNeasy Kit (Qiagen), including DNase I digestion, according to the manufacturer’s instructions. Lower RNA starting amounts, such as 25-50 μg, work very well, but higher amounts are recommended for detection of low-intensity 5’ isoforms.* *It is suggested to add spike-in which will simplify downstream data analysis (step 6 under Sequencing Data Analysis section). Usually, S. pombe cells can be added to the S. cerevisiae cells before performing RNA extraction. Alternatively, purified total S. pombe RNA can be added to the total purified S. cerevisiae RNA. Spike-in should represent ~1-1.5% of total target cells or RNA.*
5. Denature RNA sample from step 4 for 5 min at 65°C and then immediately cool on ice.
6. Pulse-spin the sample tube and mix 200 μl of denatured RNA sample with 200 μl of washed oligo d(T)25 magnetic beads from step 3. Incubate the tube for 30 min at RT at 500-550 rpm. Collect the beads on a magnetic stand and discard the supernatant.
7. Wash the beads with 500 μl of 1 × annealing buffer. Collect the beads on a magnetic stand and discard the supernatant. Repeat the wash two more times. *Resuspend the beads by gently pipetting.*
8. Collect the beads after the final wash on a magnetic stand and discard the supernatant.
9. Resuspend the beads in 100 μl TE and incubate for 3 min at 55°C.
10. Collect the beads on a magnetic stand and promptly transfer the supernatant into a new 1.5-ml microcentrifuge tube. *Prompt transfer of the eluate (as soon as the beads settle) avoids temperature drop and mRNA polyadenylated tails re-annealing to the d(T) beads.*
11. Add 100 μl of 2× annealing buffer to the supernatant from step 10.
12. Repeat mRNA enrichment steps 5—8.
13. Unite the beads from the 4 tubes by resuspending with total of 72 μl H_2_O via pipetting (18 μl each tube).
14. Incubate for 3 min at 55°C. Collect the beads on a magnetic stand and promptly transfer the eluate into a fresh 1.5-ml microcentrifuge tube.

### Fragmentation of mRNA

15. Add 8 μl fragmentation buffer (from NEBNext^®^ Magnesium RNA Fragmentation Module) to 72 μl eluate from step 14 and incubate for 3 min at 94°C. *Precise fragmentation timing is critical.*
16. Promptly transfer on ice, add 8 μl fragmentation stop buffer and vortex (or mix by tapping with fingers).
17. Add 8.8 μl 3 M sodium acetate, 1 μl 15 mg/ml GlycoBlue and 220 μl 100% ethanol to the fragmented mRNA sample and mix by inverting couple of times.
18. Flash freeze in liquid nitrogen for 5 min or keep on dry ice (–80°C) for 30 min. *This reaction can be kept overnight at –20°C.*
19. Centrifuge at 20,000 × g for 30 min at 4°C to precipitate RNA. *The small RNA pellet will co-precipitate with GlycoBlue.*
20. Discard the supernatant and gently wash the pellet with 1 ml of 70% ethanol. Centrifuge at 20,000 g for 5 min at RT. Discard all the supernatant carefully without disturbing the pellet.
21. Air-dry the pellet for 10 min at RT. Resuspend the air-dried RNA in 10 μl of nuclease-free H_2_O.
22. Denature the RNA sample for 5 min at 65°C and then immediately cool on ice. Pulse-spin the sample tube to collect the droplets.

### Dephosphorylation of 5’ and 3’ ends

23. To the RNA sample from step 22 add:

1.5 μl T4 RNA ligase buffer
2 μl SUPERase.In
24. Mix by tapping and add 2 μl Quick CIP. Mix by gently tapping the tube. Incubate for 30 min at 37°C. *In general, vortexing the samples containing an enzyme should be avoided but it is still crucial to mix the reaction. Mixing by pipetting increases handling time and chances for RNA contamination. The best practice is to mix the tube by gently taping with fingers several times.*
25. Inactivate Quick CIP for 2 min at 80°C and cool the sample on ice. Pulse-spin the sample tube to collect the droplets.

### Ligation of pre-adenylated, 3’ adapter

26. Heat 2 μl 5 μM Adapter A (Table 1) for 2 min at 70°C and cool on ice (per sample; scale appropriately). Pulse-spin the sample tube to collect the droplets.
27. To the sample from step 25, add:

2 μl Adapter A from step 26
1.5 μl T4 RNA ligase buffer
9 μl 50% PEG8000 Mix by slowly pipetting up and down. *Mind that PEG is very viscous. For multiple samples, make master mix.*
28. Add 2 μl T4 RNA ligase 2 truncated and mix by gently tapping the tube with fingers. Incubate for 2 h at 25°C and inactivate for 2 min at 70°C. Cool the sample on ice.
29. Pulse-spin the sample tube to collect the droplets. Add 2 μl 25 μM Primer B (Table 1). Mix by vortexing and pulse-spin the sample.

### Decapping of 5’ ends and ligation of 5’ adapters

30. Split reaction from the previous step in two separate tubes (each with 15.5 μl).
31. Add 2.5 μl mRNA decapping enzyme (MDE) to one tube (treat) and 2.5 μl H_2_O to the other tube (control). Mix by tapping the tubes.
32. Incubate for 1 h at 37°C, for 2 min at 70°C, and for 5 min at 37°C. *If needed, this reaction can be kept on ice until step 33 is done.*
33. Denature 1 μl 25 μM Adapter C (Table 1) for 2 min at 70°C and cool on ice for at least 1 min. Add 0.5 μl 100 mM ATP to the Adapter C (per sample; scale appropriately). *This step can be performed while RNA samples are incubated in step 32 (and kept on ice until step 32 is done).*
34. Add 1.5 μl ATP-Adapter C mix from step 33 to the sample from step 32 and mix by vortexing.
35. Add 1 μl T4 RNA ligase 1 and mix by gently taping with fingers. Incubate for 2 h at 25°C. *After ligation, the reaction can be kept overnight at –20°C.*

### Reverse transcription and library amplification

36. Add 7 μl 5× RT buffer to the sample from step 35. Incubate for 5 min at 65°C and cool on ice. Pulse spin.
37. Add following to the sample from the previous step:

3.5 μl 100 mM DTT
2.5 μl 10 mM dNTP mix
1 μl SUPERase.In Mix by vortexing and pulse-spin the sample. *For multiple samples, make master mix containing the above components. Add 7 μl master mix per sample.*
38. Add 1 μl RTIII and mix by gently taping with fingers. Incubate for 1 h at 50 °C, inactivate for 15 min at 70°C and cool down to 4°C.
39. Add PCR reaction components to the sample from step 38.

10 μl 5× Phusion HF buffer
2 μl 10 mM dNTP mix
1 μl 25 mM universal forward primer (Table 1)
1 μl 25 mM reverse index primer (Table 1)
1 μl Phusion HF DNA polymerase
40. Transfer reaction in 0.5 ml tubes and amplify using following cycle:

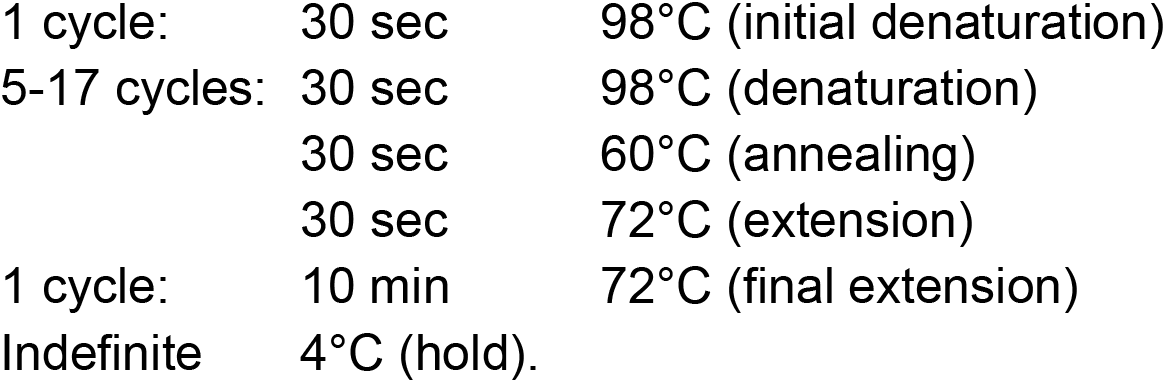 *Mind that higher number of PCR cycles may increase the sequencing duplicates.*

### Purification of amplified DNA library using beads

These steps are performed similarly as described in Jin et al., 2015 with minor modifications.

41. Equilibrate the DNA SizeSelector-I SPRI magnetic beads to RT for 20 min. *Before taking an aliquot, mix the beads thoroughly by inverting and/or swiveling the bottle.*
42. Transfer the DNA library from step 40 into a new 1.5-ml microcentrifuge tube. Add 55 μl nuclease-free H2O to adjust the final volume to 100 μl.
43. Add 200 μl of mixed SPRI bead slurry to the DNA sample from step 42 and mix by thoroughly pipetting. *Do not allow the SPRI beads to settle before adding to the samples – always mix beforehand.*
44. Incubate the mixture for 20-30 min at RT. Collect the beads on a magnetic stand and carefully discard the supernatant.
45. Wash the beads with 500 μl of freshly made 70% ethanol. Collect the beads on a magnetic stand and discard the supernatant. Repeat the wash once (total of two washes).
46. Air-dry the beads on a magnetic stand for 30 min or until the beads are completely dry. *A completely dried beads will show small “cracks” and other irregularities that are visible with careful inspection. Over-drying the beads can lead to the sample loss.*
47. Resuspend the dried beads in 22 μl of H2O and incubate for 10 min at RT. Collect the beads on a magnetic stand. Transfer only 20 μl of the supernatant into a new 1.5-ml microcentrifuge tube. The DNA library can be stored at –20°C or –80°C. *When transferring the supernatant, any carry-over of beads should be avoided. If some beads are accidentally picked up, allow the beads to re-settle on the magnet and repeat the attempt to remove the supernatant.*

### Evaluation of DNA sequencing library

48. Examine the quality and concentration of the purified DNA library using Agilent Bioanalyzer analysis (High-sensitivity DNA Kit) (Fig. 2). *The concentration of the DNA library is calculated using only those DNA fragments with sizes ranging from 175 to 800 bp (Fig. 2). A small amount of DNA smaller than 175 bp will remain in SPRI-beads-purified samples. This includes library DNAs with very short cDNA inserts, PCR primer dimers, and residual PCR primers. Gel purification is required to further extract DNA fragments with sizes only in the desired 175 to 800 bp range (Fig. 2). If multiple uniquely indexed DNA libraries will be sequenced in a single multiplexed sequencing run, the DNA libraries can be pooled according to the desired molar ratio at this stage before further gel purification. In determining the molar ratio, consider only the concentration of DNA fragments in the 175 to 800 bp range for each sample.*
49. Add an appropriate amount of 6× Gel Loading Dye to the DNA library and load the mixture on an 8% TBE polyacrylamide gel. Run at 100 V for 5 min, and then at 160V for 50 min. *Dilute PCR Marker and 100 bp DNA Ladder 20x (PCR Marker to 15 ng/μl final, 100 bp DNA Ladder to 25 ng/μl final). Load the same volume of sample and 20x diluted ladders. Use PCR Marker and 100 bp DNA Ladder to determine DNA fragments in the 175 to 800 bp range (Fig. 2).*
50. Stain the gel with SYBR Gold (1:10,000 dilution) in 40 ml TBE buffer for 3 min at RT with gentle shaking.
51. Excise DNA fragments in the 175 to 800 bp range with a disposable scalpel on a Transilluminator (Fig. 2).
52. Transfer the excised gel piece to a 0.5-ml thin-walled PCR tube with the bottom punctured with a 20-G needle. Nest the 0.5-ml PCR tube on top of a new 1.5-ml microcentrifuge tube.
53. Centrifuge the nested tubes at 20,000 × g for 5 min at RT, to effectively pulverize the relatively large gel piece into smaller pieces by squeezing the gel through the hole in the pierced tube.
54. Add 400 μl of gel extraction buffer to resuspend the pulverized gel pieces. Incubate overnight at RT with agitation (~1000-1500 rpm).
55. Transfer the whole gel/buffer mixture to the filter chamber of a Costar Spin-X centrifuge tube. Centrifuge for 20 min at 20,000 × g, RT.
56. Transfer 400 μl of the eluate from the collection tube into a new 1.5-ml microcentrifuge tube. Add 1 μl of 15 mg/ml GlycoBlue to the eluted DNA sample and mix well. Add 1000 μl of 100% ethanol, mix well, and incubate for 30 min at –80°C.
57. Centrifuge at 20,000 × g for 30 min at 4°C, to precipitate DNA.
58. Discard the supernatant and gently wash the pellet with 1 ml μl of 70% ethanol. Centrifuge at 20,000 × g for 5 min at 4°C. Discard all the supernatant carefully without disturbing the pellet.
59. Air-dry the pellet for 10 min. Resuspend the air-dried DNA in 10-20 μl of H_2_O. *The DNA sequencing library can be stored at –20°C or –80°C. The quality and concentration of the gel-purified DNA library can be examined by Agilent Bioanalyzer analysis (High-sensitivity DNA Kit) as in step 48.*
60. Paired-end sequence the library with Illumina (I used NextSeq Mid and NextSeq High) using the sequencing and index sequencing primers listed in Table 1. *Usually, 10 μl of 10 nM library is submitted for the sequencing.*

**Figure 2.**
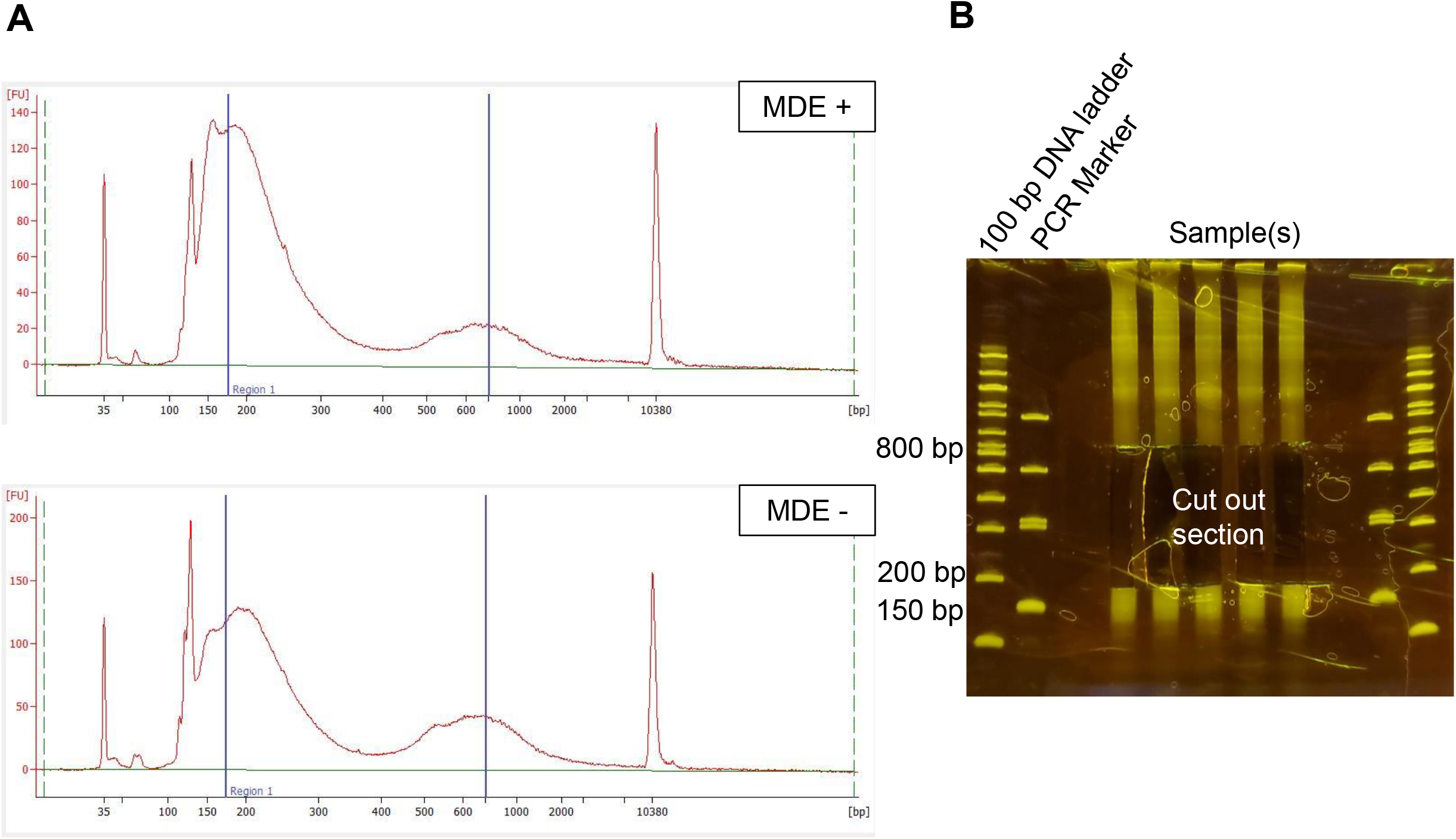
Expected results in the Bioanalyzer electrogram and acrylamide gel. A. Agilent Bioanalyzer electrogram (high-sensitivity DNA kit) of a SPRI-beads purified DNA library for MDE+and MDE-samples are shown. The DNA peaks at 35 bp and 10380 bp are high sensitivity DNA markers used in Agilent Bioanalyzer analysis. DNA fragments smaller than 175 bp include library DNAs with very short cDNA inserts, PCR primer dimers, and residual PCR primers. B. 8% TBE acrylamide gel purification is required to extract only DNA fragments between 175 bp and 800 bp for Illumina sequencing, and corresponding section of the gel is cut out. PCR Marker and 100 bp ladder indicate DNA sizes.

## SEQUENCING DATA ANALYSIS

This part is suggestion for data analysis. Both treated (MDE+) and control (MDE-) samples are processed.

1. Align paired-end sequencing reads with the following command:

bowtie2 -x genome_specific_bowtie_index –5 4 –3 16 --no-mixed --no-discordant - -dovetail -X 1000 (–5 4 –3 16 is used for 37 nt read length, –5 4 –3 55 is used for 76 nt long reads).
2. Remove duplicates from the aligned reads by considering both read pairs and the alignment location. *Truthful duplicates have the same unique identifier (random nucleotide combinations as in Adapter A and Adapter C, Table 1) and the same (or highly similar) read sequences for both read pairs at the same genomic location.*
3. Utilize pair-end read mapping information to derive genomic coordinates for the aligned fragments (usually in the form of a .bed file).
4. Obtain 5’ isoform location by trimming fragments to obtain first nt position past the 5’ adapter or the first nucleotide of the insert between the adapters (for plus and minus strand separately). *Note that removal of 4 random nucleotides from the 5’ end is performed during bowtie2 alignment and start or end of the fragments (depending on the plus or the minus strand) represent 5’ isoform location.*
5. Derive total 5’ counts per genomic coordinate separately for each strand.
6. Correct 5’ isoform reads by subtracting control per nt counts (MDE-) from the treated 5’ isoform reads (MDE+) for every nt position. *To obtain normalization (i.e. scaling) factor for treat/control, samples are aligned to the S. pombe genome as in step 3 (if S. pombe is used as spike-in). Alternatively, samples could be normalized according to the total read number. Mind that no large adjustments should be needed (scaling factors not larger than ~1.2).*
7. Individual loci can be explored in IGV (Integrated Genome Browser). To do so, sorted (normalized) 5’ isoform .bed file can be converted to .bam file (bedtools), for which .bai index can be generated (samtools).
8. To obtain effective isoform usage, isoforms can be normalized to the transcription values as follows:

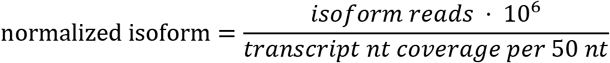

with 50 nt window being 25 nt up- and 25 nt downstream from the isoform location.
9. Using the sliding window tactics, maximum isoform within the windows can be looked for and the associated features quantified: percent signal at the most used (maximum) coordinate, maximum isoform intensities, distances between maximum isoforms, distance between the first and the last isoform, and isoform frequencies (number of distinct coordinates) (Moqtader et al., 2013). *Alone standing isoforms can be excluded from the sliding window statistics.*
10. De novo isoform-specific nucleotide frequencies can be calculated.
11. Locations of interest, such as 5’UTRs, can be preselected for the analysis.

## REAGENST AND SOLUTIONS

### Annealing buffer (100 ml), 2×

2 ml of 1 M Tris·Cl, pH 7.5 (20 mM final; RNase-free)
2.4 ml of 5 M NaCl (120 mM final; RNase-free)
400 μl of 0.5 M EDTA, pH 8 (2 mM final; RNase-free)
Adjust to 100 ml with nuclease-free H_2_O
Store at RT
*Mind that TRIS buffers require pH re-adjustments every year.*

### Gel extraction buffer

1 ml of 1 M Tris·Cl, pH 8 (10 mM final)
6 ml of 5 M NaCl (300 mM final)
200 μl of 0.5 M EDTA, pH 8 (1 mM final) Adjust to 100 ml with nuclease-free H_2_O Store at RT

### TE buffer

1 ml of 1 M Tris·Cl, pH 7.5 (10 mM final; RNase-free)
200 μl of 0.5 M EDTA,pH 8 (1 mM final; RNase-free)
Adjust to 100 ml with nuclease-free H_2_O
Store at RT

## COMMENTARY

### Background Information

Synthesis of the transcripts with alternative 5’ UTRs is a common process (Carninci et al., 2006) proposed to allow for the additional level of gene expression regulation (Davuluri et al., 2008). Changing genes’ 5’ UTR boundaries can affect mRNA translatability (Rojas-Duran and Gilbert, 2012), mRNA stability (Gu et al., 2010), and mRNA function (Davuluri et al., 2008). While single examples, such as development-, differentiation- or environment-dependent utilization of distinct 5’ transcript isoforms of the same gene have been identified (Davuluri et al., 2008, Mitschke et al., 2011, Feng et al., 2016, Kurihara et al., 2018, Lu et al., 2020, Ray et al., 2020), it can be appreciated that there are a lot of unknowns about genome-wide 5’ isoform patterns and their contribution to the gene regulation. Thus, development of simplified protocols in conjunction with decreasing next generation sequencing costs can allow for more efficient research progression in the field of 5’ isoforms.

Protocols for studying 5’ isoforms can be classified in several categories: cap-trapping, template-switching reverse transcription and oligo-capping (Machida and Lin, 2014, Policastro and Zentner, 2021). Cap-trapping (or CapSMART) involves enrichment of the capped mRNA via immunoprecipitation, or utilization of a chemical modification of the cap in conjunction with biotin enrichment. While having the advantage of providing 5’ isoform pool only, these methods still require elimination of the 5’-cap, removal of the biotin tag, fragmentation, cDNA making (as in other methods). Pre-population of capped mRNA may add to the protocol complexity compared to the other two procedures. Template-switching method involves ligation of the stop adapter to the non-capped molecules to be eliminated in the downstream steps (when the capped molecules are selected). In my hands and reported by others (Pelechano et al., 2016), this ligation step is inefficient and any molecules failing to ligate the stop adapter, including background signal which are not a representation of capped 5’ ends, would count as a positive signal. Protocols that implement oligo-capping to study 5’ transcript ends were described (Tsuchihara et al., 2009, Ni et al, 2010, Gu et al., 2012, Arribere and Gilbert, 2013, Machida and Lin, 2014, Pelechano et al., 2014, Pelechano et al., 2016). This one is different in that i) It significantly reduces the timing and handling steps of the library preparation. Thus, after mRNA enrichment and fragmentation reaction, all dephosphorylation, decapping, adapter ligation, reverse transcription, and PCR are performed in the same tube. There is no need to purify the reaction, but only add components and change the temperatures. It eliminates the need for biotinylated adapters and the usage of streptavidin beads. ii) It avoids artifacts stemming from amplicons (cDNA-DNA generation) onto which adapters are ligated. Instead, both adapters are ligated directly onto RNA. In the recent dephosphorylation-decapping methods involving indirect library construction (adapter ligation onto DNA made from RNA), only 10% of the sequenced material is usable for the 5’ isoform analysis (Pelechano et al., 2016). iii) Magnesium fragmentation is advantageously exploited to fragment RNA for adapter ligation, without generating confounding 5’ ends (i.e. the ones which would stand for capped 5’). iv) It introduces background control and allows for studying low intensity isoforms. Traditional 5’ isoform data analysis involves setting up the threshold according to which, for instance, isoforms below a certain number of reads are not counted as truthful isoforms. This approach limits the information about abundant, low intensity isoforms which are readily present throughout the whole genome (Gvozdenov et al., in preparation). In my approach, I introduced negative control – a sample identical to the treated in terms of the artifacts. Thus, background subtraction of the artifacts, traditionally not implemented for 5’ isoform studies, eliminates the need for setting a threshold and is very advantageous for identifying and studying low intensity 5’ isoforms.

### Critical Parameters and Troubleshooting

#### Starting material

To study 5’ isoform heterogeneity, it is desirable to identify less specific, low intensity isoforms that do not correspond to the frequently used ones within 5’UTR. For this reason, it is recommended to start with 1 μg mRNA (1% from the total RNA). Protocol works with lower amounts (25-50 μg starting total mRNA and less) but this may lead to the detection of primarily most used isoforms.

#### Ligations

Ligation of the 5’ adapter to RNA fragments with T4 RNA Ligase 1 can be very inefficient. Ligation for 2 h at 25°C supplemented with PEG, usage of highly concentrated T4 RNA ligase 1 (30,000 units/ml) and a good starting RNA amount greatly circumvents this problem.

Ligation with T4 RNA ligase 2 truncated in conjunction with pre-adenylated Adapter A is more efficient than ligation of Adapter C with T4 RNA ligase 1. It is crucial to ligate Adapter A to the RNA at the lower Adapter A concentration before the ligation of Adapter C at the higher concentration. Reverse order will produce a large amount of dimers on the cost of losing the target insert material. Any remaining dimers with correctly ordered ligation (Adapter A and then Adapter C) are successfully cleaned with SPRI beads and gel extraction (Jin et al., 2015).

#### Fragmentation

The fragmentation method here utilizes metal ions and heat to generate RNA fragments with 5’ hydroxyl and 3’phosphate ends (NEB). Thus, it does not produce new phosphorylated 5’. Ligation of the 5’ adapter proceeds only with phosphorylated 5’ RNA ends, which are generated once capped 5’ ends are decapped. Dephosphatase (Quick CIP) is used post fragmentation (and before decapping) to eliminate any preexisting 5’ phosphate ends, and fragmentation-generated 3’ phosphate termini. Unphosphorylated 3’ end is needed for the ligation of the pre-adenylated 3’ adapter.

Fragmentation timing is very important as too short fragmentation can lead to incomplete fragmentation and, vice versa, too long fragmentation can over-degrade RNA. Following manufacturer’s instructions (250 ng mRNA per 18 μl, NEB) and 3-4 min at 94°C is optimal to obtain 175-500 bp final library. For any variations, it is suggested to verify size distribution of the RNA fragments with Agilent Bioanalyzer analysis (RNA 6000 Pico Kit).

#### Reaction conditions and heat inactivatable enzymes

T4 RNA ligase buffer is crucial both for 5’ and 3’ adapter ligation reactions. I tested diphosphatase (Quick CIP) and mRNA decapping enzyme MDE to be very efficient in this same buffer. All enzymes are heat inactivatable. This allows for completing all dephosphorylation, ligation of 3’, decapping and ligation of 5’ (together with reverse transcription and PCR) in the same tube. Reverse transcription and PCR require addition of separate buffers, but these reactions are not impaired by the presence of the existing T4 RNA ligase buffer.

#### Control sample

Ideally, 5’ isoform signal is derived after aligning the sequencing reads to the reference genome, removing duplicates and trimming the paired-end fragments. Unlike 3’ isoforms, for which truthful isoform signal can be distinguished based on the internal polyAs- (or polyTs-) containing reads, obtaining accurate 5’ isoforms rely on the series of successful enzymatic reactions. Thus, a good control is needed to eliminate false positive signal. The control sample here is identical to the treated reaction up to the point when mRNA decapping enzyme is added. The control is the same sample during the cell growth/treatment, RNA extraction, DNase treatment, polyA enrichment, fragmentation and even 3’ adapter ligation. Only the last few steps, such as decapping (no decapping for control), 5’ ligation, RT and PCR are performed independently, i.e. the same sample is split into two reactions. This allows for the excellent control and elimination of the false positive 5’ isoform signals in the analysis.

### Understanding Results

For yeast, it is expected to obtain a very good signal if the number of successfully mapped sequencing reads and the genome length are at least in 1 to 1 ratio. A snapshot from the genome browser shows the expected 5’ isoforms for the MDE+ treated, MDE-control, and the treated after normalization (control subtraction) (Fig. 3). Subtraction helps with the elimination of the non-specific signals (Fig. 3), which is especially seen in the meta-analysis plot (Fig. 4). 5’ isoform metagene analysis with respect to *Saccharomyces cerevisiae* annotated start codons indicates 5’ isoforms peak within 5’UTR (Fig. 4). The meta-analysis was done for two different sequencing depths (2.8 and 17.7 million 5’ isoform reads after normalizing to the control sample). For the higher sequencing depth samples (17.7 million hits), control signal (MDE-) increased within the coding and 5’ region compared to the lower sequencing depth (2.8 million hits) without noticeable background. Overall, most of the non-specific, false 5’ isoforms come from the locations with higher transcriptional intensity (Fig. 3, Fig. 4).

**Figure 3.**
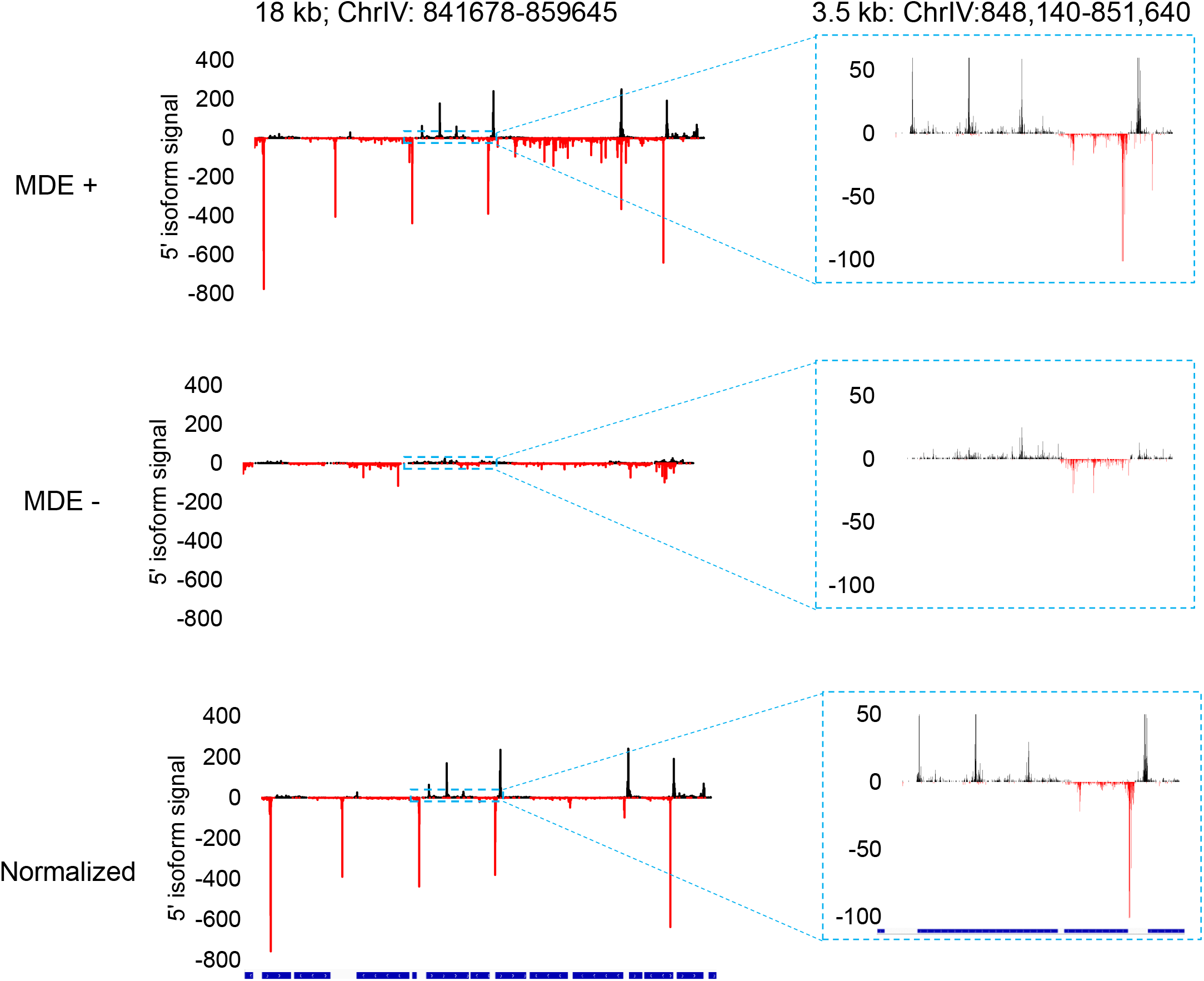
Anticipated results as in the genome browser. (Left) 5’ isoforms within 18 kb of the indicated genomic region. 5’ isoform MDE + is the treated sample (with mRNA decapping enzyme), MDE - is control (no mRNA decapping enzyme), and normalized is MDE + after normalization with MDE -. (Right) 3.5-kb section from the 18-kb window was enlarged to visualize low intensity 5’ isoforms.

**Figure 4.**
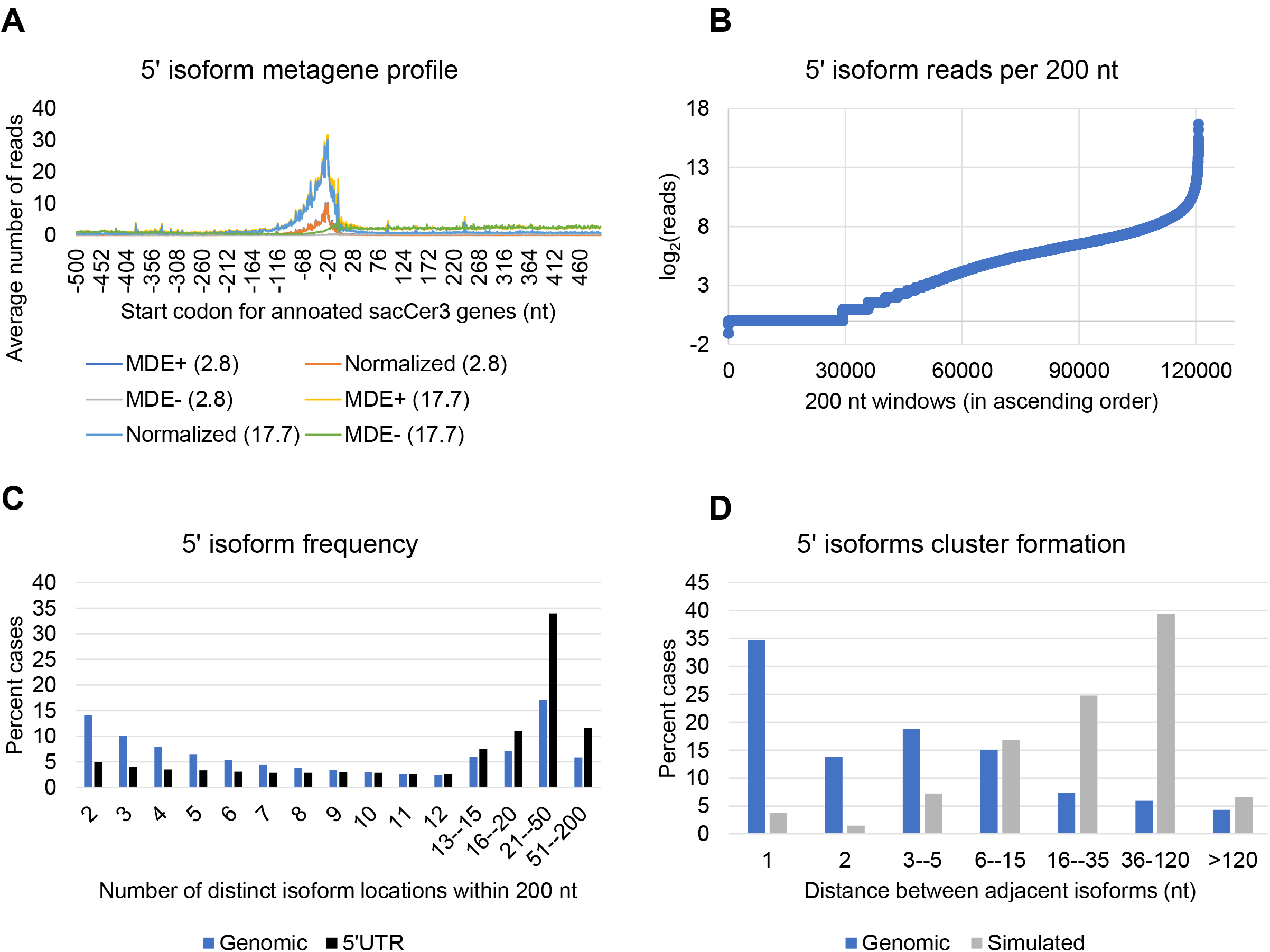
5’ isoform distributions. A. The 5’ isoform metagene profile was generated by calculating average number of isoforms with respect to the *Saccharomyces cerevisiae* start codons for annotated genes. The metagene curves were plotted for the samples with 2.8 and 17.7 million hits after normalizing all hits with the control (normalized), as well as not normalized (MDE+ only) and control samples (MDE-). B. The number of 5’ isoform reads were counted per discrete, non-overlapping 200 nt windows and plotted as log2(reads) in ascending order. C. 5’ isoform frequencies were obtained by using 200 nt sliding bins to identify and quantify the number of distinct coordinates per every bin. D. The frequency of distances between the adjacent 5’ isoforms for genomic and simulated data (truly random).

5’ isoform reads can have differential intensities and densities per genomic unit (Fig. 3), which can be seen by counting the isoform reads within genomic non-overlapping 200 nt windows (Fig. 4). For higher sequencing depth (17.7 million hits for the yeast genome after background subtraction), a quarter of the genome contains no 5’ isoforms, whereas very strong 5’ isoform occurrence is present for 1/6^th^ of the genome (log2(reads)>8) (Fig. 4). High number of 5’ isoform reads are expected within 5’UTR, whereas low and sometimes medium high 5’ isoform reads are expected for the rest of the genome (Fig. 3, Fig. 4).

A gene of interest or genome-wide analysis can be subject to other differential isoform profiles that can be examined. A sliding window tactics was utilized before to analyze 3’ isoform features (Gvozdenov et al., in preparation). Isoform frequency plot shows that 5’ UTRs contain a higher number of detectable distinct isoform locations when compared to the whole genome’s frequencies, as expected (Fig. 4). It is of note that 5’ isoforms still tend to occur in clusters as the distances between adjacent isoforms are much closer than by chance (“by chance” refers to simulated data with random isoform locations) (Fig.4). Finally, 5’ isoform motif signatures can be obtained by looking at the nucleotide signatures up- and downstream of the target 5’ isoforms (Fig. 5). These features can be important for analyzing 5’ isoform *cis* and *trans* determinants and thereby regulation of the affected genomic location by the utilization of specific 5’ isoform.

**Figure 5.**
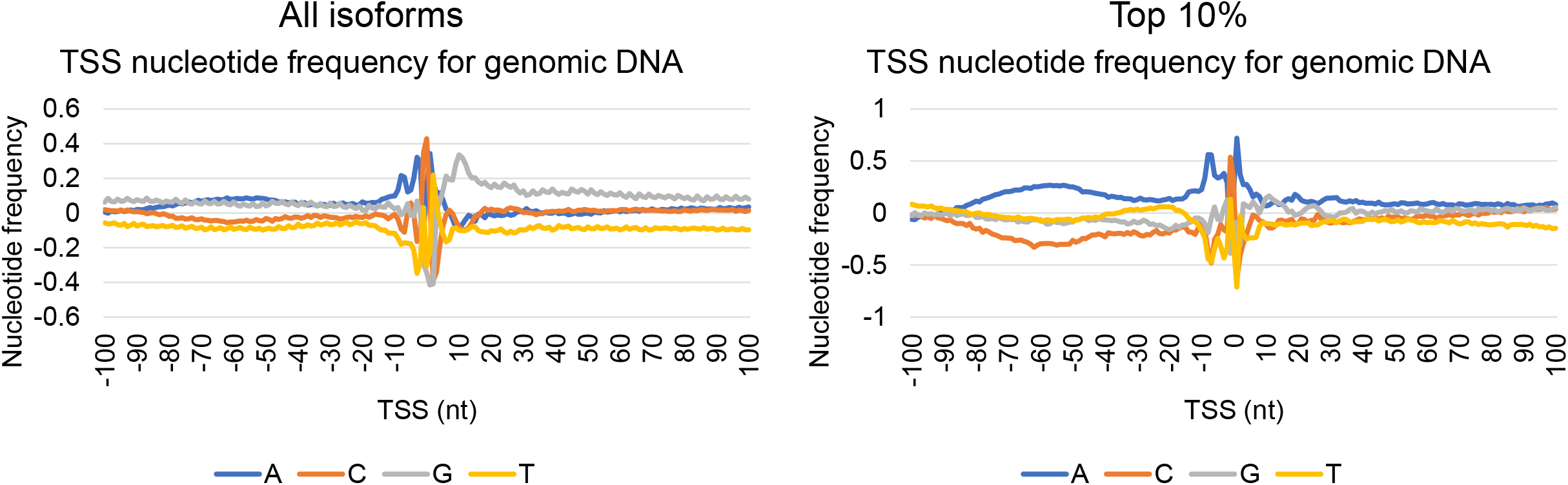
5’ isoform nucleotide signatures. Nucleotide frequencies for all normalized and top 10% 5’ isoforms (according to the isoform intensity). For each of the 100 nt up- and downstream from the isoform location, averaged nt frequencies were normalized to the occurrence of each nucleotide in the genome.

### Time consideration

The protocol described here can be completed within 3 days. 3-4 days are needed if RNA extraction is added prior to this procedure. Data analysis can be completed in 2 days to a week.

## DATA AVAILABILITY STATEMENT

Data sets are deposited with NCBI GEO under accession number GSE216450. Python scripts for data analysis are available from the author upon reasonable request.

## ACKNOWLEDGEMENTS

This project was funded by NIH research grant 1F32GM140555 to Z.G.

